# Two closed conformations of CRAF require the 14-3-3 binding motifs and cysteine-rich domain to be intact in live cells

**DOI:** 10.1101/2022.10.21.513144

**Authors:** Kenji Okamoto, Yasushi Sako

**Affiliations:** Cellular Informatics Laboratory, Cluster for Pioneering Research, RIKEN, 2-1, Hirosawa, Wako, Saitama, 351-0198, Japan

**Keywords:** protein conformation, intramolecular complex, phosphorylation, single-molecule FRET, ALEX

## Abstract

The protein rapidly accelerated fibrosarcoma (RAF) is a kinase downstream of the membrane protein RAS in the cellular signal transduction system. In the structure of RAF, the N- and C-terminus domains are connected with a flexible linker. The open/close dynamics and dimerization of RAF are thought to regulate its activity, although the details of these conformations are unknown, especially in live cells. In this work, we used alternating laser excitation to measure cytosolic CRAF in live HeLa cells and obtained single-molecule Förster resonance energy transfer (smFRET) distributions of the structural states. We compared the results for wild-type (WT)-CRAF before and after epidermal growth factor (EGF) stimulation, with mutations of the 14-3-3 binding sites and cysteine-rich domain, and an N-terminus truncation. The smFRET distributions of full-length CRAFs were analyzed by global fitting with three beta distributions. Our results suggested that a 14-3-3 dimer bound to two sites on a single CRAF molecule and induced the formation of the autoinhibitory closed conformation. There were two closed conformations, which the majority of WT-CRAF adopted. These two conformations showed different responsiveness to EGF stimulation.

## 1 Introduction

The rapidly accelerated fibrosarcoma (RAF) family of proteins has been studied extensively for decades because they play important roles in signal transduction downstream of the membrane protein RAS, and they are oncoproteins closely related to cancers (reviewed in Ref. [1]). RAF family proteins (ARAF, BRAF, and CRAF) have highly homologous amino acid sequences. RAF consists of three parts (Fig. 1A): the N-terminus, containing the RAS binding domain (RBD) and the cysteine-rich domain (CRD); a flexible linker with several phosphorylation sites; and the C-terminus, containing the catalytic domain (CAD). There are two 14-3-3 binding motifs for phosphoserine in the linker and C-terminus. CRAF has these motifs centered at serines 259 and 621, which increases the affinity with 14-3-3 when the serines are phosphorylated. The inactive form of CRAF is probably an autoinhibitory closed conformation, and the interaction between the N- and C-termini in the closed conformation suppress kinase activity [2–4]. It has been reported that a 14-3-3 dimer bridges two separate sites in a single RAF molecule to form a closed structure [5], or that 14-3-3 dimerizes different RAF molecules by connecting their kinase domains [6,7]. The CRD in the N-terminus domain may also interact with the C-terminus domain [2–4] and 14-3-3 [2,8,9]. RAF activation is initiated by the recruitment of RAF to the plasma membrane via the membrane protein RAS binding in its active GTP-form to the RBD of RAF [10,11]. After a temporary interaction with RAS, during which the CRAF structure opens [12], CRAF dissociates from the membrane [12,13]. During activation, several sites, including the N-terminal acidic motif (^338^SSYY^341^ in CRAF) or the activation loop near the kinase pocket on the CAD, are phosphorylated, whereas pS259 is dephosphorylated and 14-3-3 is detached from S259 [14,15].

**Figure 1:**
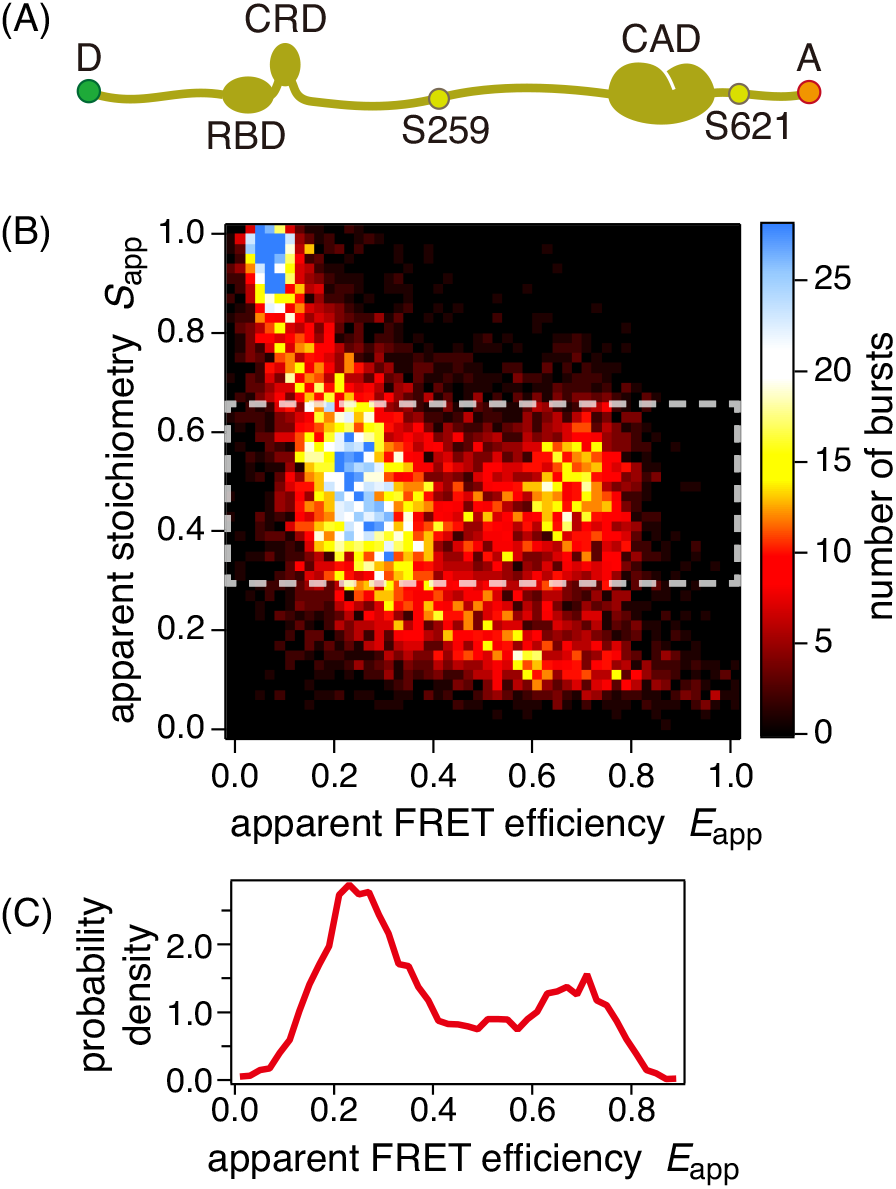
(A) Structure of CRAF. Three domains (RBD, CRD, and CAD) are connected with flexible linkers. Serines at positions 259 and 621 are 14-3-3 binding sites. Donor (D) and acceptor (A) dyes are attached at the N- and C-termini, respectively, for the FRET measurement. (B) Example of an *S*_app_ vs *E*_app_ two-dimensional histogram for CRAF_WT_ in a live HeLa cell. (C) Example of an smFRET histogram of a single cell extracted in the medium *S*_app_ range (0.3 ≤ *S*_app_ ≤ 0.65; white dashed box) of (B).

However, our understanding of the behavior of RAF in cells is still fragmentary. Furthermore, little is known about the structure of full-length RAF, although changes in this structure are believed to be important in the activation process. Information about the structure of RAF has been obtained mainly by in vitro biochemical or molecular biology experiments, which typically investigate binding interactions or interfaces, or by structural analyses such as X-ray crystallography or NMR, which give the three-dimensional structure of rigid domains. Two groups have independently reported the BRAF structure complexed with a 14-3-3 dimer in insect cells obtained by cryogenic electron microscopy (cryo-EM) [16,17]. The structure suggested intramolecular complex formation, including N- and C-termini bound on a 14-3-3 dimer before activation [16]. However, the full-length structure of RAF in live mammalian cells has not yet been reported. Förster resonance energy transfer (FRET) imaging has been used to detect the openness of the RAF structure in live cells [12,18], and some mutant RAFs may tend to have an open structure. However, observation of the ensemble average did not provide details of the molecular structures. Single-molecule imaging experiments have observed the opening of the RAF structure upon binding to RAS [12] and have analyzed the binding kinetics to RAS [13]. Because total internal reflection fluorescence microscopy captures molecules only on the plasma membrane, only RAF molecules interacting with RAS can be observed. Nevertheless, RAF is a cytosolic protein; therefore, to understand the structures of RAF in the resting and active states, it is crucial to observe the molecular behavior of cytosolic RAF.

Previously, we conducted single-molecule Förster resonance energy transfer (smFRET) measurements of cytosolic CRAFs using the alternating laser excitation (ALEX) technique, and we found that CRAFs in multiple structural states coexist in live cells [19]. In this paper, we describe further smFRET measurements of several kinds of mutant cytosolic CRAFs, especially with modifications to the interaction sites for 14-3-3 protein. We propose a structural model that consistently explains our results and supports conventional hypotheses on the RAF structure.

## 2 Results

### 2.1 Wild-type and S621A mutant CRAF

We have previously conducted ALEX measurements of single RAF molecules in the cytoplasm of HeLa cells and obtained the distributions of the apparent FRET efficiency, *E*_app_, for wild-type (WT)-CRAF (CRAF_WT_) or a mutant with a substitution from serine to alanine at 621 (CRAF_S621A_) [19]. In the present study, we extended the measurements to various RAF mutants. In the analysis of the *E*_app_ distribution, singly labeled CRAFs with one of the dyes mislabeled or photobleached were excluded owing to the apparent stoichiometry, *S*_app_. Examples of a *S*_app_ vs *E*_app_ two-dimensional histogram and an extracted *E*_app_ histogram in a single cell are shown in Fig. 1B and C, respectively.

We accumulated ALEX experimental data and obtained the average histograms from multiple cells (Fig. 2A and 2C). The results for individual cells are shown in Supplementary Fig. S1A, B, and D. Typically, the distributions for CRAF_WT_ consisted of two peaks before and after epidermal growth factor (EGF) stimulation. The variance among cells was small, as it was for all other conditions shown later, which indicated the high reproducibility of the experiments and that composition of structural states was highly regulated. EGF stimulation only caused a slight leftward shift of the left-hand peak in the *E*_app_ distribution. The slight decrease of the right-hand peak may not be important because the decrease was smaller than the variance. The shift of the left-hand peak suggested that this peak contained at least two structural states and that EGF stimulation shifted the populations between the states (Fig. 2B). Therefore, CRAF_WT_ had at least three structural states, namely, the low-FRET (LF), medium-FRET (MF), and high-FRET (HF) states. There was a non-zero distribution between the MF and LF peaks, which may have corresponded to bursts, including transitions between the LF and MF states and the HF states. CRAF_S621A_ yielded an almost single-peak distribution (Fig. 2C), which may correspond to the LF state, with a small right-hand peak corresponding to the HF state. The results for CRAF_WT_ and CRAF_S621A_ were consistent with our previous results [19].

**Figure 2:**
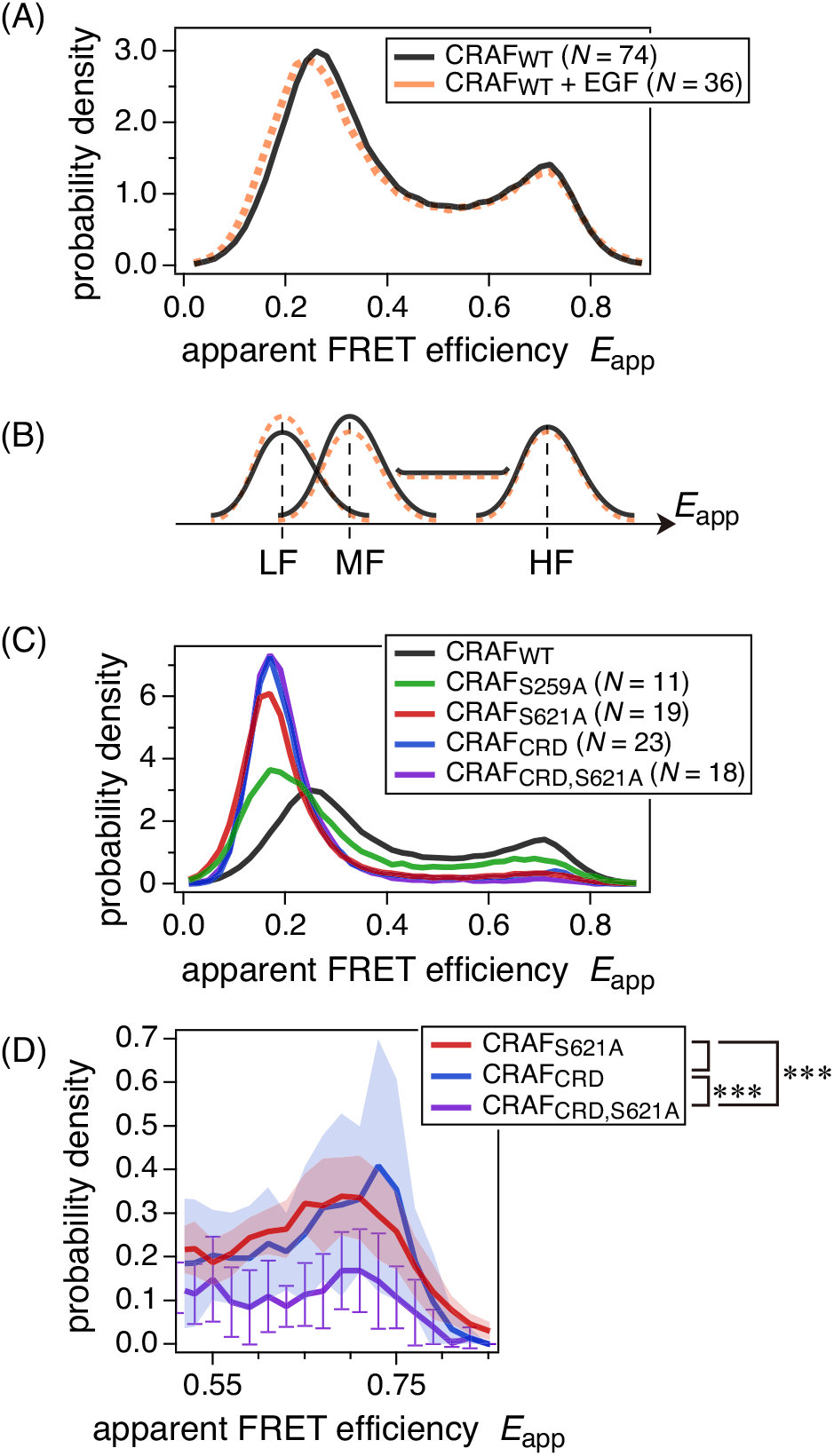
(A) Average *E*_app_ histograms for CRAF_WT_ before and after EGF stimulation. (B) Interpretation of the histograms in (A) with low-FRET (LF), medium-FRET (MF), and high-FRET (HF) peaks. The slight leftward shift of the left-hand peak may be due to a population shift between LF and MF. The flat distribution between MF and HF may be due to bursts including state transitions. (C)Average *E*_app_ histograms for CRAF_WT_, CRAF_S259A_, CRAF_S621A_, CRAF_CRD_, and CRAF_CRD,S621A_. (D) Magnification of the right-hand peak in (C). Bands and error bars represent the standard deviations among cells. Student’s *t*-test was applied against the right-hand peak heights evaluated by fitting to a beta distribution with parameters for HF (Table 1).

### 2.2 Mutants to examine the role of S259

CRAF has two 14-3-3 binding motifs at S259 and S621 [5,20]. Because the results indicated that the mutation at S621 opened the CRAF structure, 14-3-3 binding may be necessary to form the closed conformation by bridging those two sites [5].

**Table 1:** Results of global fitting analysis. *E*_*j*_ and *I*_*j*_ are globally fitted parameters representing peak position and width, respectively. The sum of fractions for a single condition (mutant) is not unity because some data points are excluded from the analysis. See details in the Materials and Methods (Section 5.5). Errors are calculated using the global fitting package of IgorPro 8.04.

|  |  | LF | MF | HF |
| --- | --- | --- | --- | --- |
| index | $j$ | 1 | 2 | 3 |
| peak parameters | $E_j$ | 0.17±0.00086 | 0.27±0.0047 | 0.70±0.0098 |
| | $I_j$ | 62.3±2.2 | 38.9±6.0 | 42.5±11 |
| peak fractions | CRAF<sub>wt</sub> | 0.10±0.029 | 0.49±0.037 | 0.24±0.033 |
|  | CRAF<sub>wt</sub> + EGF | 0.16±0.026 | 0.43±0.034 | 0.23±0.032 |
|  | CRAF<sub>S259A</sub> | 0.39±0.022 | 0.31±0.028 | 0.14±0.023 |
|  | CRAF<sub>S621A</sub> | 0.71±0.018 | 0.12±0.022 | 0.06±0.017 |
|  | CRAF<sub>CRD</sub> | 0.77±0.019 | 0.14±0.024 | 0.06±0.017 |
|  | CRAF<sub>CRD,S621A</sub> | 0.80±0.020 | 0.17±0.025 | 0.03±0.016 |

To investigate the importance of another 14-3-3 binding site at S259, we investigated full-length CRAF with an S259A mutation, CRAF_S259A_ (Fig. 2C and Supplementary Fig. S1C). The population shifted from higher FRET (MF and HF) to lower FRET (LF) structures compared with the CRAF_WT_ results, suggesting the importance of 14-3-3 in forming closed conformations.

To investigate the ability of S259 itself to bind to 14-3-3, we prepared three mutant CRAFs, CRAF_Δ1– 245_, CRAF_Δ1–245,S259A_, and CRAFΔ1–261 with the N-terminus truncated (Fig. 3A). The main difference among the mutants was the presence or absence of a phosphorylatable serine 259. Only CRAF_Δ1–245_ could have pS259 and conform to the closed structure if 14-3-3 bridged pS259 and pS621. The results of the smFRET measurement are shown in Fig. 3B and Supplementary Fig. S2. Only CRAF_Δ1–245_ showed a distinct high-FRET peak, which suggested that 14-3-3 binding to both pS259 and pS621 induced closed conformations.

**Figure 3:**
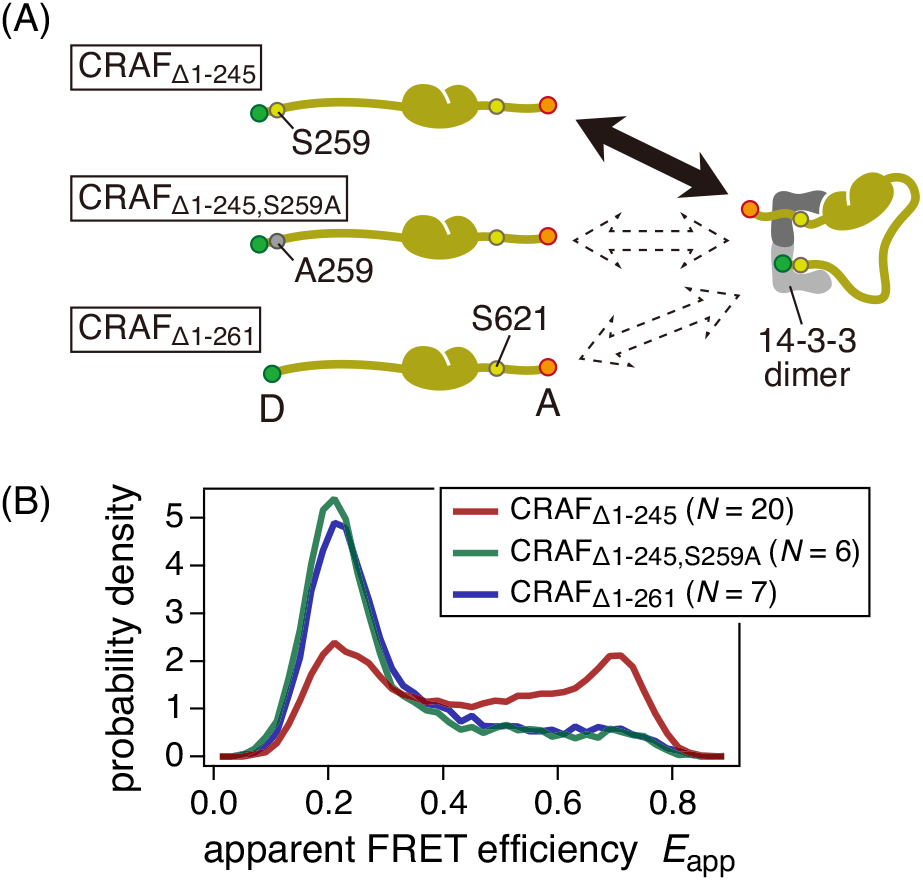
(A) N-terminus-truncated mutants of CRAF used in this study. Only CRAF_Δ1–245_ has phosphorylatable S259, whereas it is replaced with alanine in CRAF_Δ1–245,S259A_ or deleted in CRAF_Δ1– 261_. Only CRAF_Δ1–245_ may be folded by a 14-3-3 dimer. (B) Average *E*_app_ histograms of N-terminus-truncated-CRAFs.

### 2.3 Mutation in the CRD

Besides phosphorylated serines, the CRD also binds to 14-3-3 [2,8,9] and may contribute to autoinhibitory interaction between the N- and C-terminus domains [2–4]. Substitution of cysteines 165 and 168 with serines may suppress this autoinhibitory interaction [2,18]. We prepared the mutants CRAF_CRD_ and CRAF_CRD,S621A_ with a CC165, 168SS substitution. CRAF_CRD,S621A_ also had a S621A mutation.

We conducted ALEX measurements for CRAF_CRD_ and CRAF_CRD,S621A_ (Fig. 2C and D, and Supplementary Fig. S1E and F). Both yielded distributions that were almost identical to that of CRAF_S621A_. However, there was a subtle difference in results among those three mutants, in that CRAF_S621A_ and CRAF_CRD_ had a small population at the right-hand peak (HF) position, whereas CRAF_CRD,S621A_ had a smaller distribution that was insufficient to form a peak (Fig. 2D).

### 2.4 Global fitting analysis with multiple beta distributions

We obtained *E*_app_ distributions for full-length CRAFs under six different conditions (WT with/without EGF stimulation and mutants). We wanted to explain all the distributions consistently. The CRAF_WT_ distribution contained at least three states; thus, it was reasonable to assume a three-state model. The mutant CRAFs showed the same trend of the MF and the HF states decreasing while the LF state increased. We applied global fitting analysis to the *E*_app_ histograms using three beta distributions, which represent a single FRET species with an *E*_app_ peak [21,22]. However, a large number of bursts occurred between the MF and HF peaks (Fig. 2A and C), which were not included in the peaks. A major cause of these bursts was probably a structural state transition within a single burst. The bursts produced a flat distribution between peaks, which could not be expressed with a simple analytical form, preventing fitting analysis. Therefore, to exclude data points that were not included in the peaks from the fitting analysis, we used a mask (gray regions in Fig. 4A and B).

**Figure 4:**
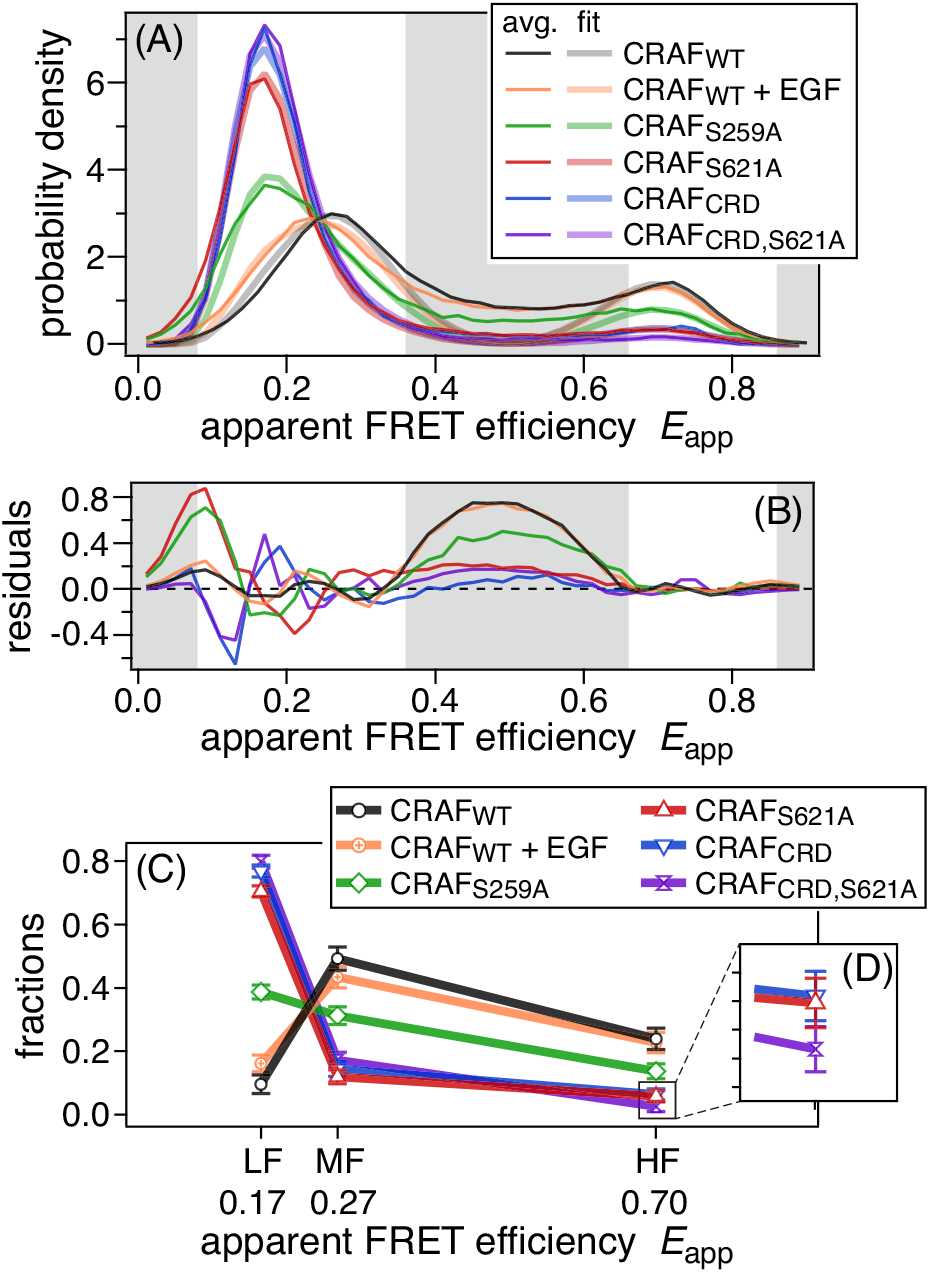
(A) Average *E*_app_ histograms (thin bright colored lines) with fitted results (thick pale colored lines). Data points in gray regions are excluded from the fitting analysis. (B) Fitting residuals. (C) Fitting results of fractions for three structural states, LF, MF, and HF. (D) Magnification of (C) for HF of CRD and S621A mutants. Error bars are fitting errors calculated by IgorPro software.

The results of global fitting are shown in Fig. 4A, and further details are shown in Fig. 6A and Supplementary Fig. S3. The experimental results (thin bright colored lines) and fitted results (thick pale colored lines) are shown together. All six histograms were fitted well with three beta distributions and the fitting residuals were suppressed, except for in the masked regions (Fig. 4B). Peak positions (apparent FRET efficiencies) and peak widths (apparent intensities) for each component and fractions of the components for each condition (mutant) are summarized in Table 1. The fractions are also shown in Fig. 4C. EGF stimulation shifted the population from the MF to the LF state and did not change the HF state substantially. Mutant CRAFs decreased the MF and HF states similarly. Both peaks were greatly decreased for CRAF_S621A_, CRAF_CRD_, and CRAF_CRD,S621A_, whereas the decrease for CRAF_S259A_ was almost half that for the other mutants. Although CRAF_S621A_, CRAF_CRD_, and CRAF_CRD,S621A_ had almost identical fractions in the fitting results, the difference in the right-hand peak (HF) in Fig. 2D was observed in the fitting results.

These fitting results do not mean that there were exactly three structural states and they represent *E*_app_ distributions. Each of the fitted peaks may include multiple structural states. If those states have slightly different FRET efficiencies, the peak appears to be broadened. Structural fluctuation with a time scale comparable to the burst duration also causes peak broadening [22]. The peak width of a single species depends on the average burst intensity, and the peak becomes narrower as the intensity increases; thus, if structural fluctuation occurs, the peak width becomes broader than that theoretically predicted with average burst intensity. In other words, fitting analysis for a broadened peak underestimates the actual average intensity. We obtained fitting results for the apparent intensities (Table 1) that were smaller than the typical average intensity of bursts (100–250), indicating that peak broadening occurred in our experiments. In addition, the data mask may not have excluded the bursts that affected the fitting analysis completely. For example, if a state transition occurred between the LF and HF states, bursts including the transition should be included in the MF peak position and may cause overestimations of the MF fraction in the low-FRET mutants (CRAF_S621A_, CRAF_CRD_, and CRAF_CRD,S621A_). Nevertheless, we believe that the FRET distribution of CRAF can be represented using three sets of structural states.

## 3 Discussion

We used the ALEX technique to conduct smFRET measurements of CRAF protein to investigate its structure in live cells. Mutations at the 14-3-3 binding sites, including CRD, S259, and S621, drastically changed the FRET distribution and shifted the population to lower FRET species. This result indicated that all sites contributed to the closed structure formation of intracellular CRAF, perhaps through interactions with 14-3-3. One possible structure for such an intramolecular complex is the cryo-EM structure obtained with full-length BRAF [16], in which N- and C-terminus domains form a complex on a 14-3-3 dimer as a scaffold (Fig. 5A). pS365 (corresponding to pS259 in CRAF) and pS729 (corresponding to pS621 in CRAF) are bound in the recognition grooves of the 14-3-3 dimer. The C-terminus CAD is tethered by pS729 and interacts with 14-3-3. The N-terminus CRD interacts with 14-3-3, the CAD, and pS365. The RBD is not clearly defined in the high-resolution cryo-EM structure, suggesting that it is dangling near CRD with a short (∼6 amino acids) linker. Because of the highly homologous amino acid sequence among RAF family proteins, CRAF may adopt a similar complex structure.

**Figure 5:**
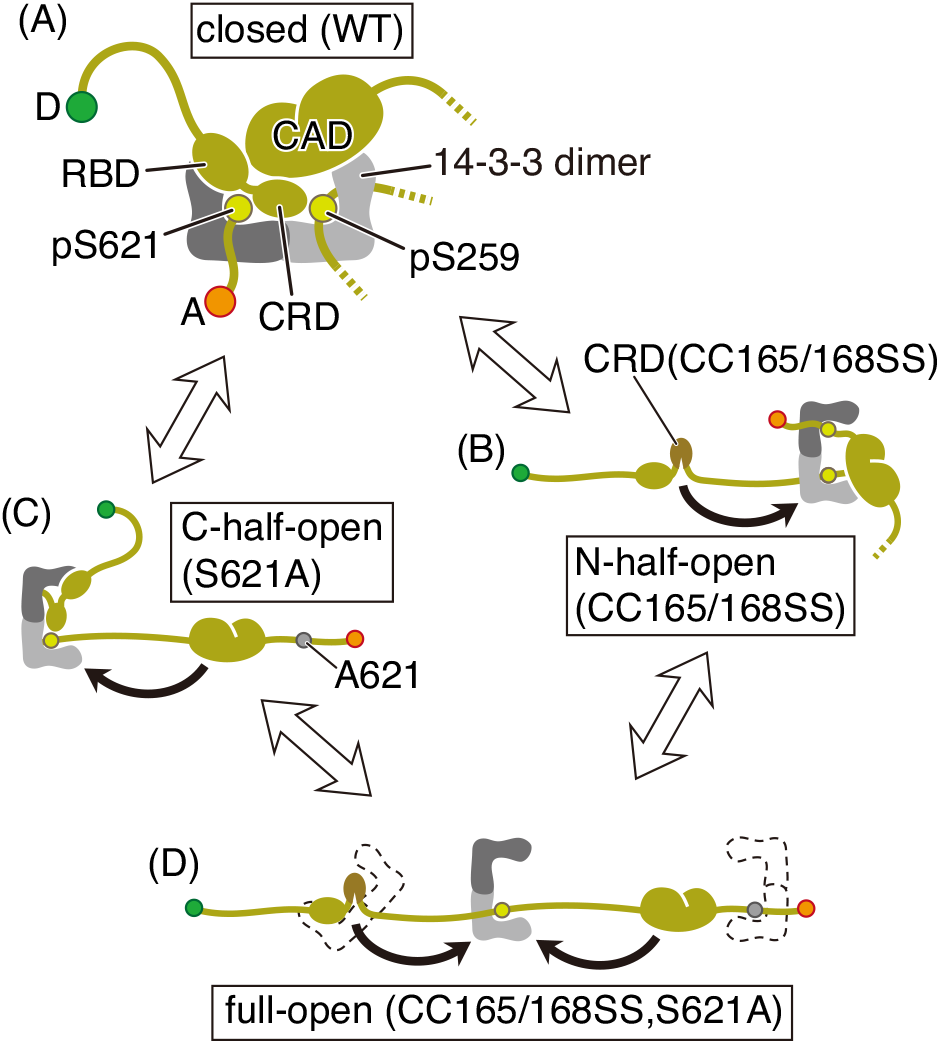
Model of conformational states of CRAF. All WT and mutant full-length CRAFs may be equilibrated among these conformations. (A) Closed conformation based on the cryo-EM structure of BRAF [16]. Both the N- and C-terminus domains are bound to a 14-3-3 dimer. A: FRET acceptor. D: FRET donor. (B) N-half-open structure with the N-terminus domain detached from 14-3-3. CRAF_CRD_ with mutations on the CRD may primarily adopt this conformation. (C) C-half-open structure with the C-terminus domain detached from 14-3-3. CRAF_S621A_ with a mutation at S621 may primarily adopt this conformation. (D) Full-open structure with both the N- and C-terminus domains detached from 14-3-3. The CRAF_CRD,S621A_ mutant may primarily adopt this conformation. A 14-3-3 dimer may bind to CRAF at a single site or be dissociated.

We assumed that there were also open conformations (Fig. 5B–D). Either the N- or C-terminus was detached from 14-3-3 in the N-half-open (Fig. 5B) or C-half-open (Fig. 5C) structure, respectively. If 14-3-3 were bound to CRAF only at a single site or were not bound to CRAF, the fully open structure may be formed (Fig. 5D). We explained the *E*_app_ distributions by assigning these structures to the LF, MF, and HF structural states. First, the closed structure, which has closely packed N- and C-termini, should show high FRET efficiency. Next, the fully open structure should show the lowest FRET efficiency among them and may be included in the LF state. CRAF_CRD_ and CRAF_S621A_ may primarily adopt the N- and C-half-open structures, respectively, because they have mutations at the 14-3-3 binding sites near the N- and C-termini, respectively. Both or either of these half-open structures could form the MF, as we assumed previously [19]. However, here we reject that hypothesis for two reasons. First, the CRAF_CRD_ and CRAF_S621A_ mutants would inhibit the interaction of either terminus with 14-3-3, but would not affect the interaction of the other terminus. If the MF state were the half-open structure, it would not be diminished by either of those mutations. Second, because the half-open structures involve the binding of S259 and 14-3-3, if the MF state were the half-open structure, it would be diminished more by the S259A mutation. Therefore, all the half- and fully open structures were included in the LF state, suggesting that both the MF and HF states were the closed conformations. However, because the MF and HF states had separate FRET peaks, their structures were different. Both structures involved interactions with both the N- and C-termini because their populations were lost in CRAF_CRD_ and CRAF_S621A_ mutants. Either structure could be similar to the cryo-EM structure (Fig. 5A), but it is unknown what the other structure was and which of the two structures corresponded to the MF and HF states. We assigned the MF and HF states as the closed conformations and the LF state as the open conformations (Fig. 6).

**Figure 6:**
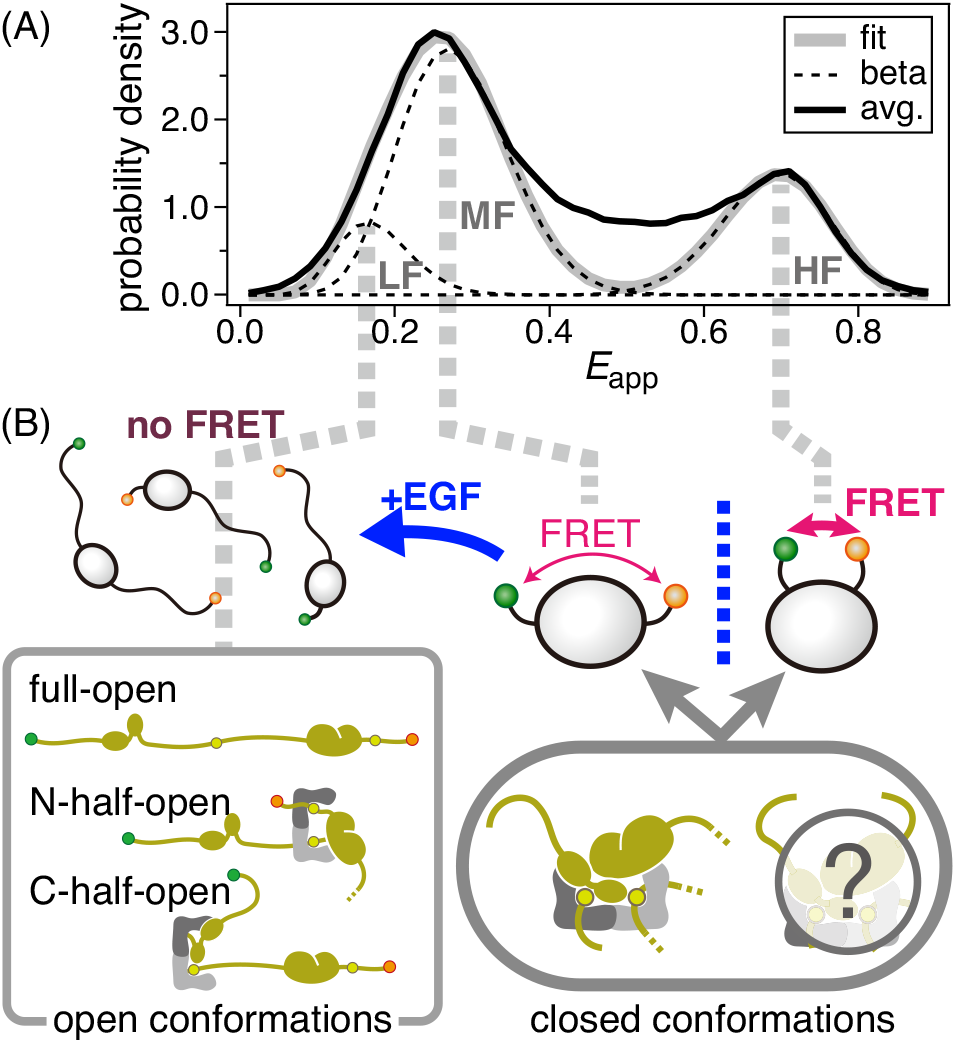
(A) Results of fitting analysis for CRAF_WT_. The average *E*_app_ histogram (black solid line), fitted result (gray solid line), and three fitted beta distributions (black dashed lines). Each of the LF, MF, and HF states is represented by a single beta distribution. (B) Speculative assignment of structural states. LF is the open conformations, including N- and C-half-open and fully open structures. MF and HF are different closed conformations. One is probably similar to the cryo-EM structure of BRAF (Figure 5A), but the other structure is unknown. See details in the Discussion section.

According to the above assignment of the states, the LF state contained the half- and fully open conformations, which should have different interdye distances. Therefore, the end-to-end distance of the CRAF molecule was too long for FRET, even in the half-open structures. The detected FRET of the LF state (*E*_app_ ∼ 0.17) was attributed to the spectral bleed-through of donor fluorescence to the acceptor detection channel, which was estimated as ∼0.07 from the position of the peak near the top-left corner in Fig. 1B and direct excitation of the acceptor by donor excitation light.

The assignments represented the static *E*_app_ distributions as the result of the equilibrium among the conformational states, which may explain the differences in the *E*_app_ distributions of CRAF_S621A_, CRAF_CRD_, and CRAF_CRD,S621A_. The distributions appeared similar, but the right-hand peaks corresponding to the HF state were small for CRAF_S621A_ and CRAF_CRD_, whereas the peak was absent for CRAF_CRD,S621A_ (Fig. 2C and D). These differences were also present in the fitting results (Fig. 4C and D and Table 1). For example, CRAF_S621A_ primarily adopted the C-half open conformation, but the interaction between the C-terminus and 14-3-3 still occurred with decreased affinity, which resulted in the small HF peak. A similar interaction may occur at the N-terminus for the CRAF_CRD_ mutant. However, for CRAF_CRD,S621A_, which may primarily adopt the fully open conformation, the closed conformation may be hardly formed because the interactions of both termini with decreased affinity must occur at the same time.

The *E*_app_ distribution for the S259A mutant showed the MF and HF peaks decreased by half compared with those for CRAF_WT_. The loss of binding between S259 and 14-3-3 did not explain the result because the FRET efficiency appeared to depend directly on the binding states of N- and C-termini to 14-3-3, but not directly on S259 binding. Furthermore, nearly half of CRAF_Δ1–245_, which had a phosphorylatable S259, formed a low FRET peak (Fig. 3B). These results appeared inconsistent with the hypothetical model that the phosphorylation state is uniformly regulated and decides the structural state. Consequently, we performed phosphorylation assays by using western blotting and mass spectroscopy analyses (Supplementary Results S2.1). The western blotting results showed that the phosphorylation of S259 and S621 was decreased in the mutants, except for pS259 of the S621A mutant (Supplementary Fig. S4B and C). The lower phosphorylation level of S621 in CRAF_S259A_ may have decreased the MF and HF peaks and some CRAF_S259A_ may have adopted the C-half-open structure. Similarly, some CRAF_CRD_ may adopt the fully open structure because of the decrease of phosphorylation at S259 and S621. The mass spectroscopy results showed relatively low phosphorylation at S259 (∼16%) in CRAF_Δ1–245_ (Supplementary Tables S1–3 and Supplementary Fig. S5), which explained the formation of the low-FRET peak (Fig. 3B). However, the phosphorylation appeared to be too low to form the high-FRET peak that involved almost half the population. The result may indicate that S259 can interact with 14-3-3 even without phosphorylation with low affinity. We examined the response of CRAF to EGF stimulation (Figs 2A and 4C and Table 1). There was little difference in the *E*_app_ distribution before and after EGF stimulation, suggesting that most CRAFs did not show a structural change in response to EGF stimulation. Because RAF activation accompanies a structural change to the open structure, most CRAFs did not respond to EGF stimulation under our experimental conditions. Only part of the MF state (6%–7% of the total CRAF) responded and changed to the LF state, as previously suggested [23]. This result suggested that the MF and HF states, which were different closed conformations, had different responsiveness to EGF stimulation or that a responsive substate is included in the MF peak in addition to the unresponsive components in the MF and HF states.

Our structural model is consistent with conventional hypotheses about the full-length structure of CRAF. For example, CRAF contains two 14-3-3 binding motifs, centered at S259 and S621 [24,25]. CRD also interacts with 14-3-3 [2,8,9]. These interactions are included in our model. In quiescent cells, CRAF is in an inactive state, in which the interaction between the N-terminus domain and CAD autoinhibits kinase activation [2–4]. Our model is consistent with such interaction because N- and C-termini are closely placed on the 14-3-3 scaffold. Although the mechanism of autoinhibition remains unclear, the interface for kinase dimerization, which is necessary for activation [6,26,27], may be obstructed by 14-3-3 [16]. Phosphorylation of S259 also suppresses catalytic activity [4,5,8,28–30]. This observation is explained by our finding that pS259 stabilizes closed conformations.

It has been proposed that 14-3-3 binds the C-termini of two different RAFs to form a kinase dimer [4,6]. In 2019, two groups, including the group that reported the closed structure (Fig. 5A), independently reported cryo-EM structures of a kinase dimer using full-length BRAF [16,17]. The kinase dimers were assumed to be the activated form because they were detected only without phosphorylation at S365 (corresponding to S259 of CRAF), which is believed to be dephosphorylated in the activation process. Because the closed structure was obtained using WT-BRAF with phosphorylated S365 [16], CRAF may also prefer to form a similar closed structure if S259 can be phosphorylated. If such kinase dimers are formed for CRAF, they may be included in the LF state, to which a part of the MF component moved after EGF stimulation. Although the kinase dimer model does not assume anything about the position of the N-terminus, the kinase dimer probably adopts the open conformation because the N-terminus was not included in the cryo-EM structure. In addition, the kinase dimer model does not explain why the S259A mutation decreased the populations of the MF and HF states. Therefore, it is highly probable that the major components of CRAFs in quiescent cells are two types of closed conformation with all CRD, S259, and S621 bound on 14-3-3.

## 4 Conclusion

The full-length structure of RAF in live cells has long been undetermined. In this paper, we measured smFRET of WT and mutant CRAF molecules in live HeLa cells. Although we did not directly observe 14-3-3 binding to RAF, we confirmed that mutation on the 14-3-3 binding motifs or CRD opened the CRAF structure, strongly suggesting that 14-3-3 binds to S259, S621, and CRD in a single CRAF molecule. These results are consistent with the previous hypotheses that 14-3-3 bridges two phosphorylated serines at 259 and 621, the N- and C-terminus domains interact to autoinhibit kinase activity, and CRD is the possible binding site for 14-3-3 on the N-terminus domain. Our findings support the conventional view that CRAF activation is initiated by the interaction of RAS with a monomeric CRAF, invalidating the autoinhibition by the interaction between the N- and C-terminus domains. Our results also suggested that WT-CRAF has two distinct closed conformations with different responsiveness to EGF stimulation, although the details of these structures remain unknown. In addition, there are still many questions, for example, about the structure, activation process, and dimerization of RAF proteins, and further investigation is required.5

## 5 Materials and Methods

### 5.1 Plasmid construction for wild-type and mutant CRAFs

Protocols for the preparation of the cDNA encoding EGFP-CRAF-HaloTag have been described previously [19]. In brief, point mutations were introduced to cDNA by site-directed mutagenesis using the QuikChange Lightning Site-Directed Mutagenesis Kit (Agilent Technologies, Inc.) and PrimeSTAR Max DNA Polymerase (Takara Bio Inc., Japan), and all mutations were confirmed by DNA sequencing.

Eight plasmids were constructed for this study, and CRAF_WT_ for WT-CRAF and CRAF_S621A_ for S621A mutant CRAFs were the same as those used in a previous study [19]. All other mutants were modified from CRAF_WT_. CRAF_S259A_ had a mutation from serine to alanine at position 259, where phosphorylated serine is a 14-3-3 binding target. CRAF_CRD_ had a mutation from two cysteines to two serines at positions 165 and 168. Because both cysteines form zinc fingers close to the N- and C-termini of the CRD [31,32], the CRD structure may have been disrupted in this mutant. CRAF_CRD,S621A_ contained both CC165/168SS and S621A mutations. CRAF_Δ1–245_, CRAF_Δ1–245,S259A_, and CRAF_Δ1–261_ were N-terminus-truncated mutants with the amino acids at positions 245 and 261 deleted from the N-terminus. CRAF_Δ1–245,S259A_ also contained the S259A mutation. GFP and HaloTag were encoded at the N- and C-termini, respectively, in all constructs for FRET measurements.

### 5.2 Cell culture and CRAF expression in live cells

HeLa cells were maintained in Dulbecco’s minimum essential medium (high glucose) using L-glutamine (FUJIFILM Wako Chemicals Co., Ltd., Japan) supplemented with 10% fetal bovine serum. In all experiments, cells transiently expressing proteins were used.

Subconfluent cells, which were at ∼10% confluence for the ALEX experiments and at less than 50% confluence for SDS-PAGE, were transfected with the EGFP-CRAF-HaloTag expression vector using Lipofectamine3000 (Thermo Fisher Scientific Inc.). Concentrations of cDNA and P3000 reagent were reduced to 10%–20% of those for the standard protocol and incubation time was shortened to 3 h from 2–4 days recommended for the standard protocol, to suppress expression levels of the labeled proteins. Cells were starved overnight in Eagle’s minimum essential medium (EMEM; Nissui Pharmaceutical Co., Ltd., Japan) with 1% bovine serum albumin (BSA). The HaloTag was labeled with a ligand covalently attached to the fluorescent dye tetramethylrhodamine (Promega Corp.) before measurement. Cells were kept in EMEM + 1% BSA with 15 mM HEPES (pH 7.4) (Nacalai Tesque, Inc., Japan) during measurements at 26 °C. For the +EGF conditions, EGF was added to a final concentration of 100 ng mL^−1^ after ALEX measurements were conducted on cells expressing CRAF_WT_.

### 5.3 ALEX

The apparatus and procedures for the ALEX measurements have been described previously [19]. In brief, fluorescence bursts were detected from single CRAF molecules with two-color fluorescence labels diffusing in the cytosol. The detected photons were classified into four types with two-color excitation and two detection channels, yielding four photon counts for each burst. After background subtraction, four fluorescence intensities, *I*_X→Y_, where X and Y are the donor (D) or acceptor (A) dye and represent the excitation light and detector channel, respectively, were obtained.

The apparent FRET efficiency *E*_app_ and the apparent stoichiometry *S*_app_ are calculated as

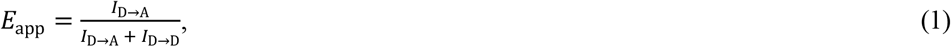

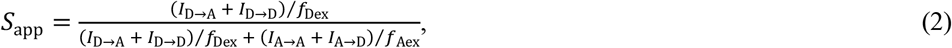

where *f*_Dex_ and *f*_Aex_ are the ALEX excitation fractions for donor and acceptor excitation, respectively. *f*_Dex_ and *f*_Aex_ were typically ∼0.7–0.8 and ∼0.2–0.3, respectively, with *f*_Dex_ + *f*_Aex_ kept slightly less than unity to avoid overlapping excitations. *E*_app_ histograms were constructed from bursts within a certain *S*_app_ range, which was adjusted for each cell (Fig. 1B and C).

We treated *E*_app_ and *S*_app_ without compensating for experimental factors, such as spectral bleed-through, direct excitation of the acceptor dye with donor excitation light, and the differences in quantum yields or detection efficiencies between dyes. The *E*_app_ distribution could be expressed by the beta distributions, enabling fitting analysis as described below, whereas the compensated histograms could not be expressed by a simple analytical form [33].

### 5.4 Global fitting of FRET histograms with beta distributions

The *E*_app_ histogram of a single species can be represented by a beta distribution [21,22]. The fitting function for a single *j*-th component is defined as

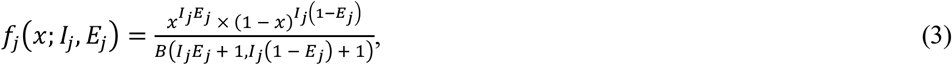

where *I*_*j*_ is the apparent intensity and *E*_*j*_ is the apparent FRET efficiency for the *j*-th component, and 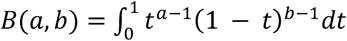 is the beta function. *E*_*j*_ represents the peak position, corresponding to the molecular structure. *I*_*j*_ determines the peak width and corresponds to the average burst intensity if the sample keeps a constant FRET efficiency. However, in actual experiments, peak width is broadened for various reasons [22]. Thus, in the analyses in this paper, *I*_*j*_ was a parameter representing peak width, and a higher *I*_*j*_ indicated a narrower peak. Different components can have different *I*_*j*_ values, even in the same measurement.

The whole *E*_app_ histogram for the *i*-th condition (mutant) is fitted with the composite distribution function,

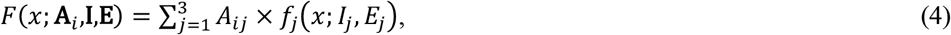

where **A**_*i*_ = {*A*_*i*1_, *A*_*i*2_, *A*_*i*3_} represents the fraction of components, **I** = {*I*_1_, *I*_2_, *I*_3_} and **E** = {*E*_1_, *E*_2_, *E*_3_}. Eq. 4 did not fit the whole *E*_app_ histograms well because there were some bursts that could not be represented by beta distributions. For example, if a state transition occurs within a burst, *E*_app_ of the burst appears somewhere between *E*_app_ of the states before and after the transition [33]. It is likely that the flat distribution in the center region of our results was caused by such bursts. Because the distribution of these bursts cannot be expressed analytically, they prevent fitting analysis. To avoid the bursts, we used a mask so that only data points contained in peaks (*E*_app_ of 0.08–0.38 and 0.66–0.88) were used for fitting analysis. Data points in the masked regions (gray regions in Fig. 4A and B) were not used in the analysis.

Six *E*_app_ histograms were fitted globally with Eq. 4 using the global fitting package of IgorPro 8.04 (WaveMetrics Inc., Lake Oswego, OR, U.S.). *I*_*j*_ and *E*_*j*_ are global fitting parameters for the *j*-th component. **A**_*i*_ was fitted to each of six conditions. Summation 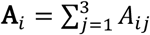 was not equal to unity due to the exclusion of some data points by masking.

## Acknowledgments

We are grateful to the Support Unit for Bio-Material Analysis, RIKEN CBS Research Resources Division, for technical help with mass spectroscopy analysis.

