## Supplementary Information for "Two closed conformations of CRAF require the 14-3-3 binding motifs and cysteine-rich domain to be intact in live cells"

### S1 Supplementary Materials and Methods

#### S1.1 SDS-PAGE and western blotting

Cells were transfected, starved, washed with Hanks' balanced salt solution, and then harvested. For the WT + EGF conditions, the cells were treated with 100 ng $mL^{-1}$ EGF (Upstate, Merck Millipore) for 10 min before washing. The cells were solubilized with SDS sample buffer, separated on 8% polyacrylamide gel, and transferred onto a PVDF membrane (Merck Millipore). The membrane was incubated with a primary antibody against GFP (Takara Bio Inc., Japan) and a secondary antibody conjugated with alkaline phosphatase (Vector Laboratories) and was stained using 5-bromo-4-chloro-3-indolyl phosphate/*p*-nitroblue tetrazolium chloride color development substrate (Promega). The primary antibody anti-Ser(P)259 RAF (Cell Signaling Technology) was used to detect phosphorylation at S259 of CRAF.

#### S1.2 Mass spectroscopy

$CRAF_{WT}$ and $CRAF_{\Delta1–245}$ were purified by immunoprecipitation using GFP-Trap agarose beads (#GTA-10, Proteintech Group, Inc.) from cell lysate. Electrophoresed proteins were stained with CBB and single bands were obtained. Each gel band was excised and cut into small pieces. After washing and destaining the gel pieces, cysteine residues were reduced by dithiothreitol and alkylated with iodoacetamide. The proteins were digested with modified trypsin, and then the resulting peptides were subjected to liquid chromatography (LC)–tandem mass spectroscopy (MS/MS).

LC–MS/MS analysis was performed using a liquid chromatography system (EASY-nLC 1000, Thermo Fisher Scientific) coupled with a mass spectrometer (Q Exactive, Thermo Fisher Scientific) equipped with a nanospray ion source. The peptides were separated with a NANO-HPLC capillary C18 column (Nikkyo Technos, Japan; 0.075 × 150 mm, 3 μm) using the following 60 min gradient at a flow rate of 300 nL/min (solvent A: 0.1% formic acid; solvent B: 100% $CH_3CN$/0.1% formic acid): 5%–35% B over 48 min, and then 35%–65% B over 12 min. The resulting mass spectroscopy (MS) and MS/MS data were searched against the Swiss-Prot database using Proteome Discoverer (Thermo Fisher Scientific) with MASCOT search engine software (Matrix Science, UK).

### S2 Supplementary Results

#### S2.1 Estimation of phosphorylation

First, we estimated the phosphorylation degrees of the serines in the 14-3-3 binding motifs by western blotting analysis of CRAF mutants with GFP and Halo-tag at the N- and C-termini, respectively. Representative data for three independent experiments are shown in Supplementary Figure S4. The cells were treated with 100 ng $mL^{-1}$ EGF for 10 min under WT + EGF conditions. The expression of the CRAF mutants was evaluated by staining with anti-GFP. Anti-pS259 and anti-pS621 were used to evaluate phosphorylation at S259 and S621, respectively. $CRAF_{S259A}$, $CRAF_{\Delta 1–245,S259A}$, and $CRAF_{\Delta 1–261}$ lacked phosphorylatable S259. $CRAF_{S621A}$ and $CRAF_{CRD,S621A}$ mutants lacked phosphorylatable S621.

S259 phosphorylation was detected only in Δ1–256 mutants among the N-terminus-truncated mutants (Fig. S4A, dashed rectangle). The degrees of phosphorylation at S259 and S621 were evaluated as the staining intensities of anti-pS259 and anti-pS621 normalized to the protein expression (anti-GFP) (Figs S4B and C). Mutations replacing a serine with an alanine at position 259 or 621 almost eliminated the corresponding band. Unexpectedly, we found that the phosphorylation of serines also decreased in mutants in which a serine was not replaced with an alanine, except for pS259 of the S621A mutant. In particular, the decrease in phosphorylation at S259 for the CRD and Δ1–245 mutations was significant. The low phosphorylation of the Δ1–245 mutant (~3% of $CRAF_{WT}$) seemed inconsistent with ALEX results (Figs 3B and S2A), in which nearly half of $CRAF_{\Delta 1–245}$ formed a high-FRET peak, suggesting that the closed structure was formed, and hence S259 was phosphorylated in those high-FRET CRAFs.

We suspected that the inconsistency between the western blotting and ALEX results could be explained by the low affinity of anti-pS259 for $CRAF_{\Delta 1–245}$ because of the truncation of the N-terminus near S259. Therefore, we conducted MS analysis to quantify the phosphorylation at S259.

To compare the phosphorylation degrees between the $CRAF_{WT}$ and Δ1–245 CRAFs, we prepared only $CRAF_{WT}$ (A) and $CRAF_{\Delta 1–245}$ (B) for MS analysis. We conducted three independent sets of experiments, including cell preparation, transient transfection, and LC–MS/MS measurements.

S259 was detected only in peptides at the 257–275 positions in the original CRAF. The peptide [257–275] was detected with and without phosphorylation. Phosphorylation on the peptide was detected only at S259. Abundances of the peptides without (*np*) and with (*p*) phosphorylation are shown in Supplementary Table S1. The normalized abundances, which were defined as the abundances divided by the average of each, for *np* and *p* are shown in Supplementary Fig. S5A.

The three sets of measurements for A ($CRAF_{WT}$) and B ($CRAF_{\Delta 1-245}$) showed that *p* was larger than *np* for A and vice versa for B. However, the normalized abundance cannot be compared directly between *np* and *p* because the detection efficiency for *p* was probably decreased greatly by the addition of the negative charge of a phosphoryl group. In addition, cell numbers and protein expression were not uniform across the six measurements (three each for A and B), making it difficult to compare phosphorylation degrees quantitatively.

Therefore, we calculated the relative abundances, which were normalized to the total expression of the CRAF protein. Thus, the expression level of the protein was determined for each measurement. Peptides from other parts of CRAF were required as the reference for the expression level because the abundance of peptides should be proportional to total CRAF expression [1]. The reference peptide had to satisfy the following three conditions: 1) it must be from amino acid positions after 246 in the original CRAF because it must be included both in $CRAF_{WT}$ and $CRAF_{\Delta 1–245}$; 2) it must be detected uniformly without variations in length and positions caused by digestion; and 3) variations in modification, such as phosphorylation, must not be detected. We found five candidate reference peptides satisfying all the conditions at positions 276–282 (*r*1), 415–431 (*r*2), 451–462 (*r*3), 555–563 (*r*4), and 564–572 (*r*5). The abundances and normalized abundances are shown in Supplementary Table S1 and Fig. S5B, respectively. Four of the five candidate peptides showed similar trends; only the results for *r*2 were different. The relative abundances, which are *np* and *p* divided by *r*1–5, respectively, are shown in Supplementary Fig. S5C–F. We selected *r*1, *r*3, *r*4, and *r*5 as reference peptides for CRAF expression.

The mean and the standard deviation of the relative abundances were calculated for $CRAF_{WT}$ and $CRAF_{\Delta 1–245}$ without (*NP*) and with (*P*) phosphorylation from twelve relative abundances (4 references × 3 independent measurements; Figs S5D and F and Table S2). We used the equations $NP + \alpha P = \beta$ (= const.) for $CRAF_{WT}$ and $CRAF_{\Delta 1–245}$, where $\alpha$ is the coefficient that represents the difference in the detection efficiencies. By solving the simultaneous equations, $\alpha = 0.346$ and $\beta = 1.29$ were obtained and fractions of *NP* and *P* for $CRAF_{WT}$ and $CRAF_{\Delta 1–245}$, respectively, were obtained (Table S3 and Fig. S5G).

The estimated proportion of phosphorylation for $CRAF_{\Delta 1–245}$ (~16%) still appeared to contradict the ALEX results that suggested that nearly half of $CRAF_{\Delta 1–245}$ had phosphorylated S259. These results may indicate that S259 can interact with 14-3-3, even without phosphorylation with low affinity.

### Supplementary Tables

| | position | A1 | B1 | A2 | B2 | A3 | B3 |
|---|---|---|---|---|---|---|---|
| *np* | [257–275] | $2.96\times10^{7}$ | $9.28\times10^{8}$ | $1.27\times10^{8}$ | $3.39\times10^{8}$ | $1.38\times10^{8}$ | $2.57\times10^{8}$ |
| *p* | [257–275]+P | $1.22\times10^{8}$ | $1.72\times10^{8}$ | $2.36\times10^{8}$ | $5.35\times10^{7}$ | $2.89\times10^{8}$ | $3.99\times10^{7}$ |
| *r*1 | [276–282] | $5.51\times10^{8}$ | $3.43\times10^{9}$ | $9.94\times10^{8}$ | $1.10\times10^{9}$ | $1.13\times10^{9}$ | $7.07\times10^{8}$ |
| *r*2 | [415–431] | $4.41\times10^{5}$ | $1.28\times10^{7}$ | $2.36\times10^{6}$ | $8.65\times10^{6}$ | $1.76\times10^{7}$ | $3.58\times10^{7}$ |
| *r*3 | [451-462] | $7.65\times10^{7}$ | $5.58\times10^{8}$ | $1.78\times10^{8}$ | $1.47\times10^{8}$ | $2.34\times10^{8}$ | $1.26\times10^{8}$ |
| *r*4 | [555–563] | $1.23\times10^{8}$ | $5.43\times10^{8}$ | $1.43\times10^{8}$ | $2.41\times10^{8}$ | $3.30\times10^{8}$ | $2.28\times10^{8}$ |
| *r*5 | [564–572] | $4.02\times10^{8}$ | $1.87\times10^{9}$ | $5.06\times10^{8}$ | $4.83\times10^{8}$ | $4.90\times10^{8}$ | $3.48\times10^{8}$ |

Supplementary Table S1: Abundances of seven peptides detected by LC–MS/MS measurements. The amino acid positions are for the original CRAF. Three independent measurements were conducted for A ($CRAF_{WT}$) and B ($CRAF_{\Delta 1–245}$). S259 was detected only in *np* and *p*. S259 was phosphorylated in *p*. *r*1–5 are candidates for the reference peptides for the CRAF expression level.

| | $CRAF_{WT}$ | $CRAF_{\Delta 1\text{-}245}$ |
|---|---|---|
| *NP* | 0.35±0.15 | 1.08±0.20 |
| *P* | 2.70±0.57 | 0.59±0.10 |

Supplementary Table S2: Mean and standard deviations of the relative abundances, which are the ratios of the normalized abundances of the target peptide (*np* and *p*) to those of the reference peptides (*r*1–5), for $CRAF_{WT}$ and $CRAF_{\Delta 1–245}$ without (*NP*) and with (*P*) phosphorylation at S259. Twelve relative abundances (four references (*r*1, *r*3–*r*5) × three measurements) each were used for the calculation.

| | $CRAF_{WT}$ | $CRAF_{\Delta 1\text{-}245}$ |
|---|---|---|
| *NP* | 0.27±0.12 | 0.84±0.16 |
| *P* | 0.73±0.15 | 0.16±0.03 |

Supplementary Table S3: Fractions of CRAF without (*NP*) and with (*P*) phosphorylation at S259 for $CRAF_{WT}$ and $CRAF_{\Delta 1–245}$.

### Supplementary Figures

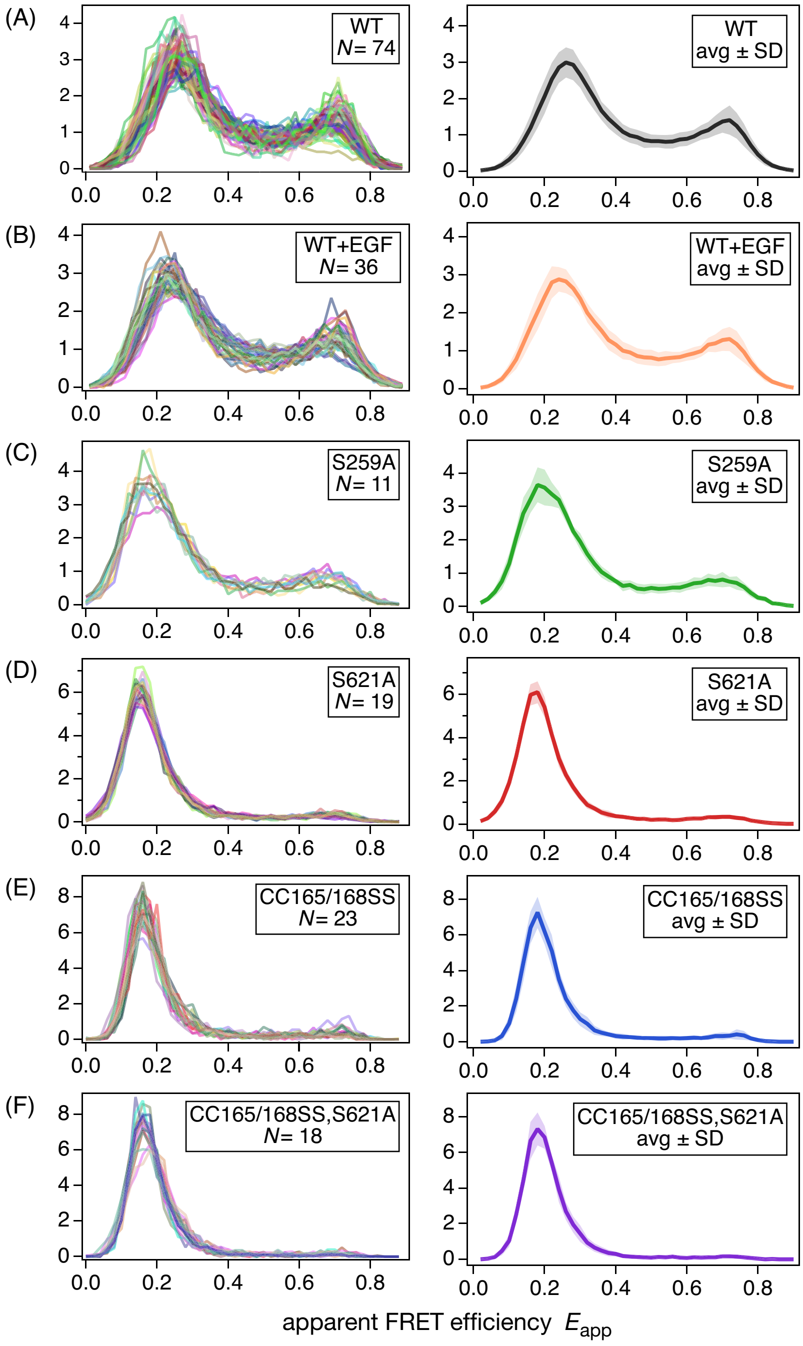


Supplementary Figure S1: $E_{app}$ histograms of full-length CRAFs under various conditions. (A) $CRAF_{WT}$, (B) $CRAF_{WT}$ after EGF stimulation, (C) $CRAF_{S259A}$, (D) $CRAF_{S621A}$, (E) $CRAF_{CRD}$, and (F) $CRAF_{CRD,S621A}$. Left-hand panels show $E_{app}$ histograms of all cells used in analyses, randomly colored to distinguish individual cells. Right-hand panels show the average $E_{app}$ histograms with bands representing the standard deviations.

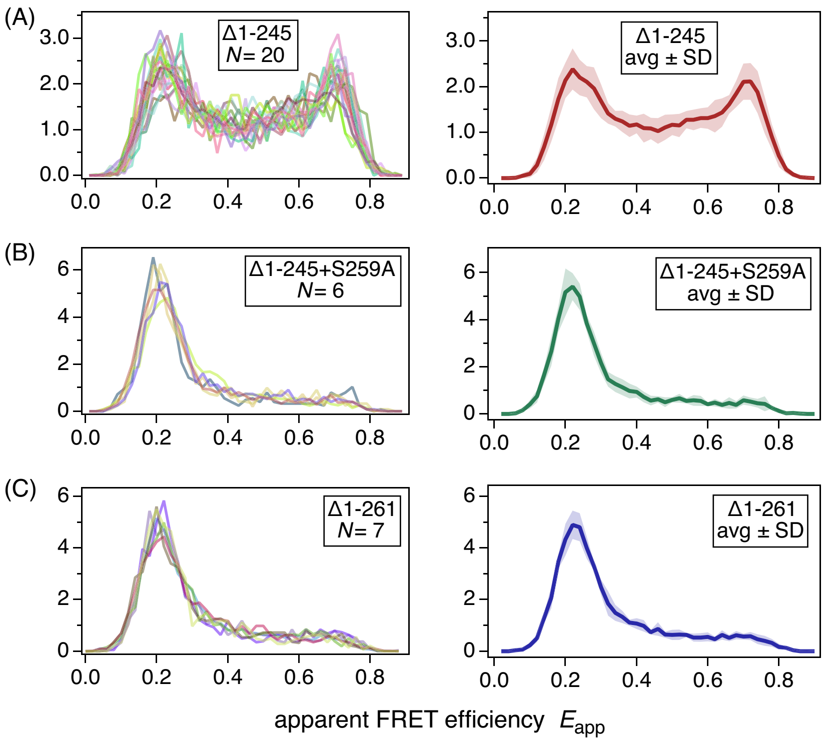


Supplementary Figure S2: $E_{app}$ histograms of N-terminus-truncated CRAFs. (A) $CRAF_{\Delta 1–245}$, (B) $CRAF_{\Delta 1–245,S259A}$, and (C) $CRAF_{\Delta 1–261}$. Left-hand panels show $E_{app}$ histograms of all cells used in the analyses, randomly colored to distinguish individual cells. Right-hand panels show the average $E_{app}$ histograms, with bands representing the standard deviations.

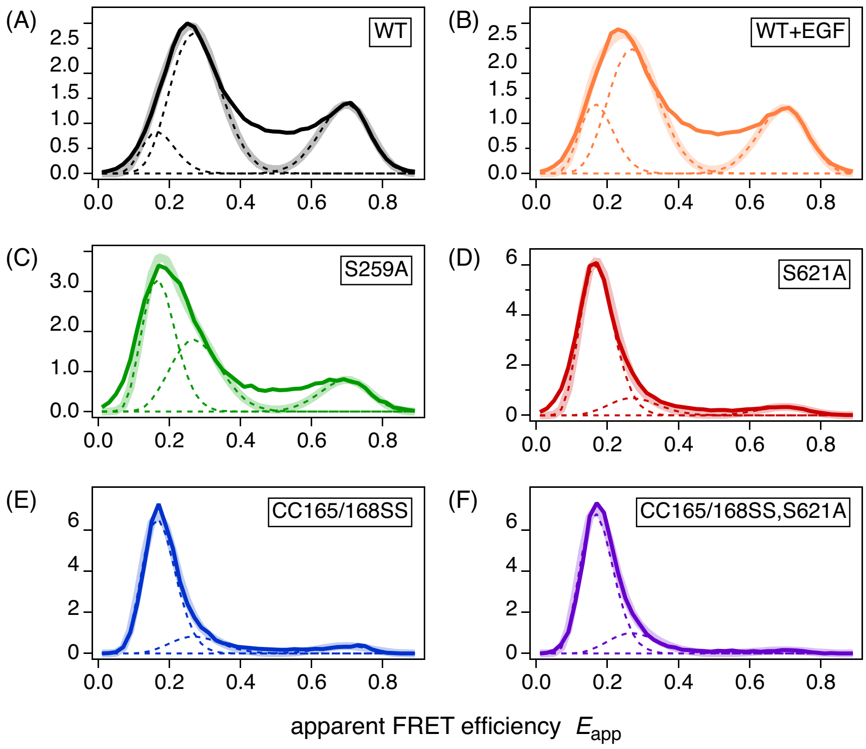


Supplementary Figure S3: Results of global fitting analysis. (A) $CRAF_{WT}$, (B) $CRAF_{WT}$ after EGF stimulation, (C) $CRAF_{S259A}$, (D) $CRAF_{S621A}$, (E) $CRAF_{CRD}$, and (F) $CRAF_{CRD,S621A}$. Thin bright colored solid lines are the average $E_{app}$ histograms, thick pale colored solid lines are the fitted results (Eq. 4), and dashed lines represent the beta distributions of the individual components (LF, MF, and HF). The peak position and width of each component are the same for all conditions (mutants).

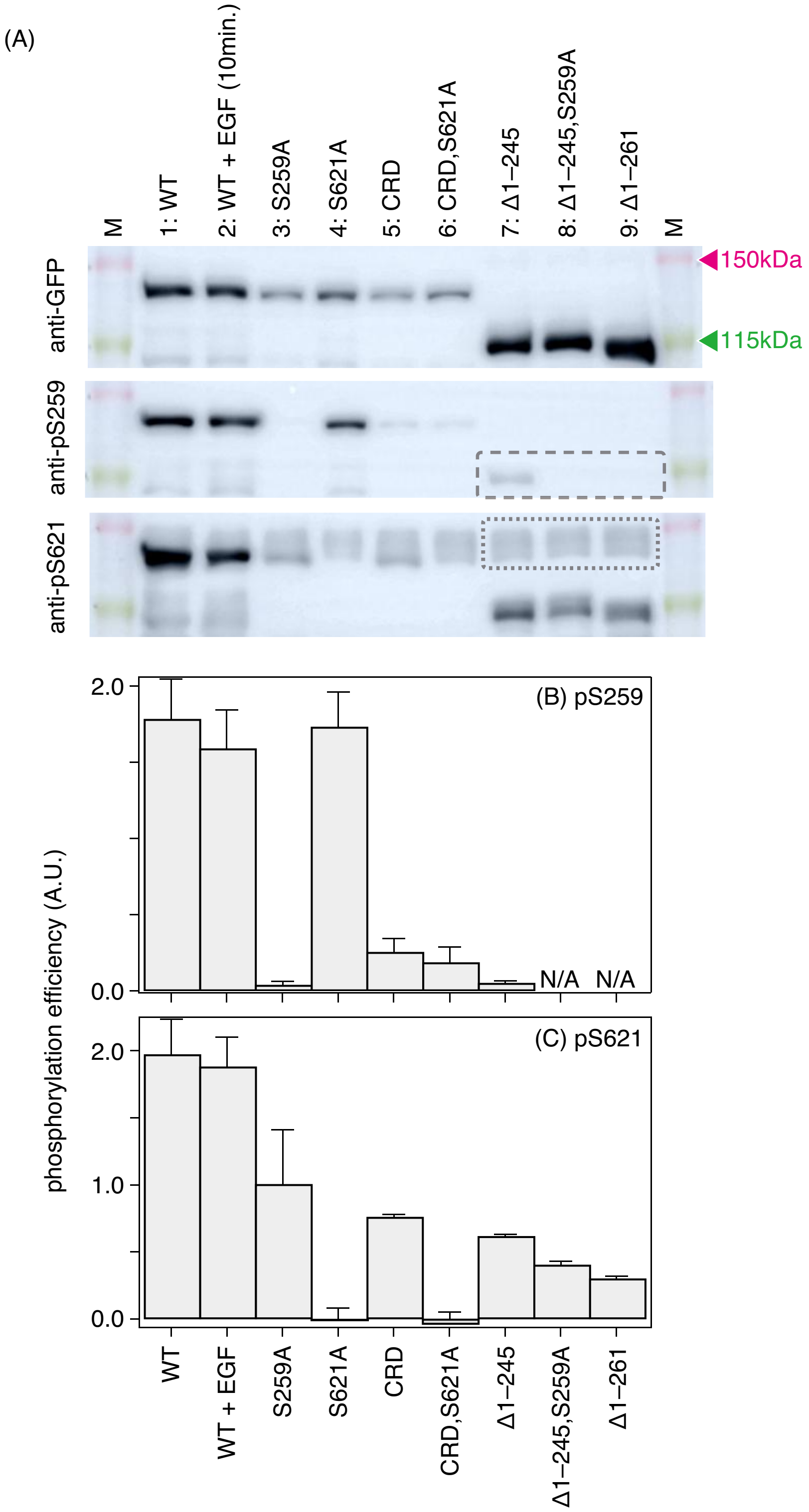


Supplementary Figure S4: (A) Western blot analysis of the phosphorylation of CRAF mutants with GFP and Halo-tag at the N- and C-termini, respectively. The cells were treated with 100 ng $mL^{-1}$ EGF for 10 min under the WT + EGF conditions. The expression of CRAF mutants was evaluated by staining with anti-GFP. Anti-pS259 and anti-pS621 were used to evaluate phosphorylation at S259 and S621, respectively. M: molecular weight marker. Representative data for three independent experiments are shown. S259 phosphorylation was detected only in Δ1–256 mutants among the N-terminus-truncated mutants (dashed rectangle). (B) Degrees of phosphorylation at S259 evaluated as the staining intensities of anti-pS259 normalized to the protein expression (anti-GFP). pS259 bands were undetectable for 8: Δ1–245, S259A, and 9: Δ1–261 mutants. (C) Degrees of phosphorylation at S621 evaluated as the staining intensities of anti-pS621 normalized to the protein expression (anti-GFP). Background intensity, which was calculated as the average of three bands in N-terminus-truncated mutant lanes (dotted rectangle), was subtracted from the anti-pS621 intensities of full-length CRAF mutants. Error bars represent the mean ± standard deviations of three independent experiments.

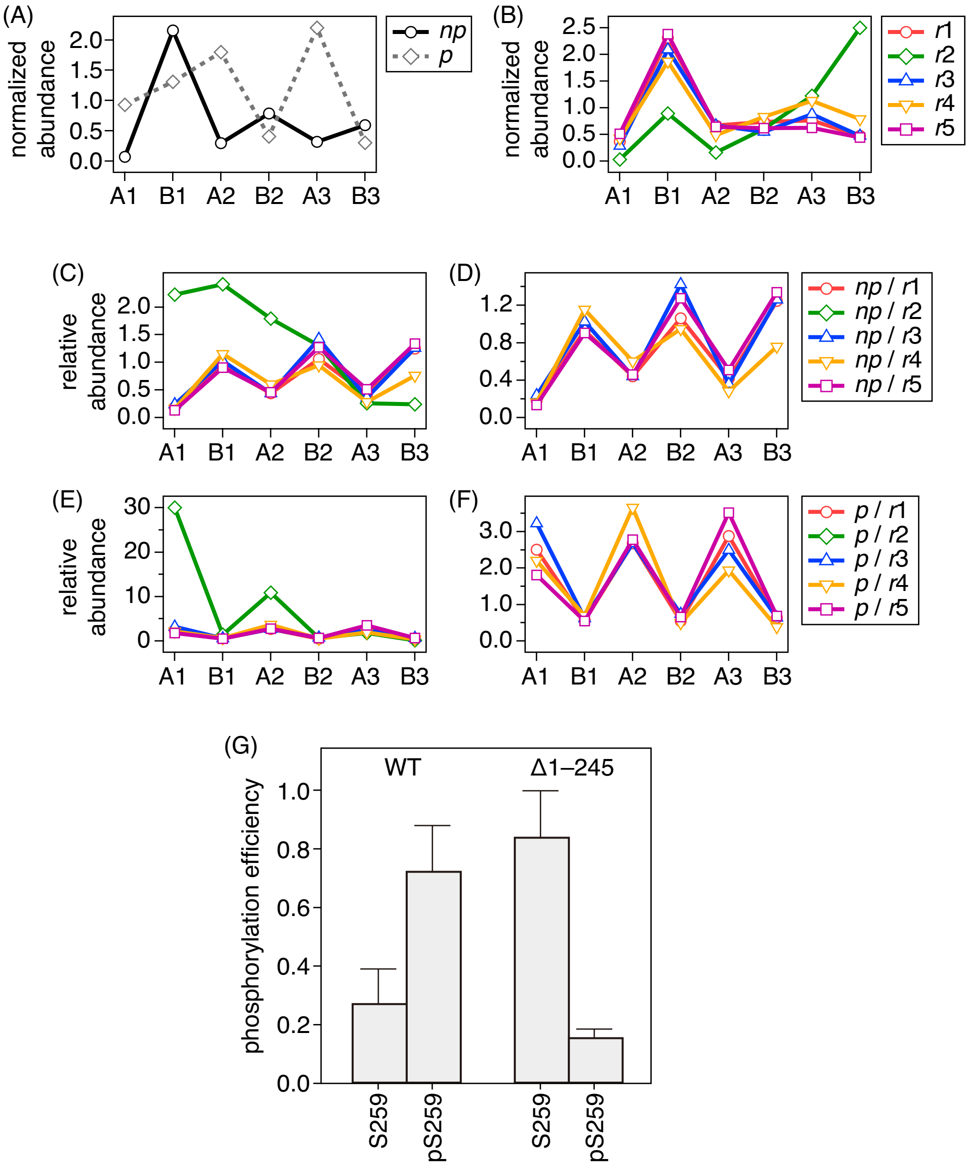


Supplementary Figure S5: LC–MS/MS results. (A) Normalized abundances of the peptides including S259 without (*np*) and with (*p*) phosphorylation at S259. (B) Normalized abundances of five candidate peptides as the reference for CRAF expression. (C) Relative abundances of the *np* peptide normalized by the reference peptides *r*1–5. (D) Same as (C), but the results for *r*2 were excluded. (E) Relative abundances of the *p* peptide normalized by reference peptides *r*1–5. (F) Same as (E), but the results for *r*2 were excluded. (G) Fractions of CRAFs with or without phosphorylation at S259 for $\mathrm{CRAF_{WT}}$ and $\mathrm{CRAF_{\Delta 1\text{–}245}}$ (Supplementary Table S3).
